# Kazal-type serine protease inhibitors from *Arabidopsis thaliana* and *Toxoplasma gondii* exhibit antimicrobial activity against plant pathogens

**DOI:** 10.1101/2025.11.28.691165

**Authors:** Manuel A. Sánchez, Karen N. Strack, Luisa F. Mendoza-Morales, Victor A. Ramos-Duarte, Eliane Perez, Valeria A. Sander, Melina Laguía-Becher, Fernando L. Pieckenstain, Marina Clemente

## Abstract

Kazal-type serine protease inhibitors (KPIs) interfere with microbial proteases and have been associated with antimicrobial activity, yet their specific role in plant protection remains poorly understood. In this work, we evaluated the antimicrobial potential of recombinant *Arabidopsis thaliana* KPI (rAtKPI-1T), recombinant *Toxoplasma gondii* KPI (rTgPI-1), and two truncated versions of rTgPI-1 (rTgPI-1_NT_ and rTgPI-1_CT_). Both rAtKPI-1T and rTgPI-1 exhibited inhibitory effects against *Pseudomonas syringae* DC3000, *P. syringae* (AvRpm1), and *P. viridiflava* in a concentration-dependent manner, with significant inhibition at 3.5 μM. The lower MIC_50_ values obtained for rTgPI-1 and its truncated forms compared to rAtKPI-1T suggest a higher antibacterial potency. Binding and immunofluorescence assays further revealed that rAtKPI-1T, rTgPI-1, and rTgPI-1_NT_ associated with bacterial surfaces, while rTgPI-1_CT_ displayed weaker or transient interactions. In addition to their antibacterial activity, rTgPI-1_NT_ and rTgPI-1_CT_ also inhibited the germination of *Botrytis cinerea* conidia. Both truncated proteins significantly reduced germination after 6 and 9 hours of incubation, with rTgPI-1_CT_ exerting a markedly stronger antifungal effect than rTgPI-1_NT_. These findings suggest that specific Kazal-type domains, particularly those in the C-terminal region, could play a critical role in suppressing early infection processes of necrotrophic fungi. Overall, this study demonstrates that rKPIs display distinct and complementary antimicrobial profiles mediated by microbial binding and protease inhibition. The contrasting activities of rTgPI-1_NT_ and rTgPI-1_CT_ highlight the value of domain-level analysis to discern functional contributions within multidomain KPIs. These results expand current knowledge on plant-derived protease inhibitors and underscore their potential as biotechnological tools for sustainable crop protection.

## Introduction

Serine protease inhibitors (SPIs) from plants are typically small proteins with a low molecular weight, detected mainly in storage tissues, such as tubers and seeds, and in the aerial parts of the plants [1]. Plant SPIs constitute one of the most abundant protein classes (between 1 and 10% of total proteins), being able to inhibit different serine proteases [2]. Several reports demonstrated that plant SPIs could participate in defense mechanisms in response to injury or attack by insects or pathogenic microorganisms (bacteria and/or fungi) [3]. In addition, plant SPIs were involved in processes related to the mobilization of reserve proteins, regulation of endogenous enzymatic activities, modulation of apoptosis or programmed cell death processes, and stabilization of defense proteins and compounds against animals, insects, and microorganisms [3,4]. Although most studies on plant SPIs have primarily focused on expanding knowledge [5], recent evidence indicates that findings on the role of ISPs may have significant implications for developing biotechnological tools to control pests and diseases caused by phytopathogens [6]. In this context, the necrotrophic fungus *Botrytis cinerea* represents one of the most relevant phytopathogens worldwide, being responsible for gray mold disease in more than 1,600 plant species, and it is widely used as a model for pathogenicity studies due to its broad host range and secretion of proteases that facilitate host tissue colonization[7] [8].

The Kazal-type SPI family (KPIs) is one of the 18 best-characterized families of SPIs and plays a regulatory role in processes involving serine proteases such as trypsin, chymotrypsin, thrombin, elastase, and/or subtilisin [9]. KPIs have a conserved motif in their amino acid sequence known as the Kazal domain [10]. More than 100 KPIs have been identified and characterized in mammals, birds, insects, parasites, fungi, and bacteria [11], demonstrating an important role in normal cell function and physiological processes [12,13]. In addition, KPIs were also linked to biological functions such as the modulation of the immune response [14–16], blood coagulation [17,18], regulation of the inflammatory response [19], and the pathogenesis of mammalian parasites and pathogenic fungi [20–23]. Furthermore, several KPIs also exhibit microorganism-binding activity by inhibiting the activity of microbial serine proteases, and show important antimicrobial activity against a wide variety of bacteria and fungi [24–26].

KPIs have also been identified in plants [27,28]; however, little is known about their specific role [29]. Previously, we identified two putative KPIs from *Arabidopsis thaliana*, which we named AtKPI-1 and AtKPI-2, that have a single Kazal-type domain [30]. Based on sequence analysis, we classified these KPIs as non-typical and non-classical Kazal-type domains with two additional cysteines to the six cysteines identified in the typical Kazal-type domains. The recombinant protein for AtKPI-1 (rAtKPI-1T) displayed high specificity for elastase and subtilisin and high inhibitory activity against the germination of *Botrytis cinerea* conidia [30]. On the other hand, some studies have shown that SPIs from non-plant organisms can efficiently inhibit serine proteases from plant pathogens [31]. On this basis, in a previous work, we analyzed the antimicrobial properties of the recombinant TgPI-1 (rTgPI-1), a non-classical KPI from *Toxoplasma gondii* with four Kazal-type domains and a potent inhibitor of trypsin, chymotrypsin, and elastase [20]. In this way, we demonstrated that similarity to rAtKPI-1T, rTgPI-1 is an effective inhibitor of *B. cinerea* conidial germination [30].

The Pseudomonadaceae family is composed of Gram-negative bacteria, with several species known to be pathogenic on a wide range of plants, including trees, vegetables, and fruits [32]. Among them, *P. syringae* stands out as a major pathogen of crops, resulting in considerable economic losses worldwide. Beyond its role as a plant pathogen, *P. syringae* can also act as an opportunistic pathogen in humans, potentially leading to infections, especially in individuals with weakened immune systems, such as immunocompromised patients [33]. Considering that, many plant SPIs have been shown to have relevant antimicrobial activity [34], and that several KPIs by inhibiting the activity of bacterial proteases also exhibit microorganism binding activity on a wide variety of bacteria and fungi [24–26,35]; in this work, we evaluated the antimicrobial activity of rAtKPI-1T and rTgPI-1 to contribute to improving the current methods for plant protection against pathogens. Also, we performed a binding assay to analyze the microbial binding activity and an indirect immunofluorescence assay (IIF) to visualize the binding of the rKPIs to live bacterial cells. Likewise, we included two truncated versions of rTgPI-1 (rTgPI-1_NT_ and rTgPI-1_CT_) to determine which domains of this protein contribute to binding activity. Thus, we demonstrated the strong potential of these rKPIs as new bacterial growth inhibitors, especially against *Pseudomonas* species and *B. cinerea*.

## Materials and methods

### Strains and culture medium

*Pseudomonas syringae* DC 3000, *P. syringae* (AvRpm1), and *P. viridiflava* were cultured on King’s medium B [36] at 28 °C for 24 h, supplemented with 50 μg/ml rifampicin. Before each experiment, one colony was used to inoculate 5 mL of liquid King’s B medium supplemented with 50 μg/ml of rifampicin, followed by overnight incubation at 28°C. Bacterial growth was monitored based on the optical density at 600 nm using a Zeltec 5000 spectrophotometer.

Conidia from *B. cinerea* strain B05.10 colonies cultivated on potato dextrose agar (PDA) plates were obtained as described by Pariani et al. [30]. Conidia were collected using sterile water containing 0.02% (v/v) Tween-20, filtered, and quantified with a hemocytometer. The suspension was adjusted to a final concentration of 1.5 × 10^4^ conidia per milliliter.

### Expression and purification of recombinant KSPIs

AtKPI-1_23-96_ (rAtKPI-1T) was expressed in *E. coli* BL21 (DE3) as described by Pariani et al. [30], while TgPI-1_24-318_ (rTgPI-1) and the truncated versions TgPI-1_24-218_ (rTgPI-1_NT_) and TgPI-1_164-318_ (rTgPI-1_CT_) were expressed in *E. coli* M15 as described in Bruno et al. [37]. All purification procedures were carried out as described previously [30,38]. Briefly, soluble non-denaturing recombinant proteins were purified using a nitrilotriacetic acid-Ni^2+^ column (Qiagen, California, USA) and eluted stepwise with 250 mM imidazole in lysis buffer. Protein concentration was measured using the Bradford assay [39]. Recombinant proteins were stored at 4 °C.

### Bacterial growth inhibition assay

The bacteriostatic activity of rATKPI-1T, rTgPI-1, and the rTgPI-1_NT_ and rTgPI-1_CT_ domains was evaluated following the method described by Donpudsa et al. [24]. Each assay was performed in duplicate for each bacterial strain. Overnight bacterial cultures were diluted 1:100 and incubated with shaking in the presence of 0, 3.5 and 7 µM of each recombinant protein. Incubation was carried out at 28 °C, and bacterial growth was monitored by measuring optical density at 600 nm for 16 hours.

### Minimum inhibitory concentration (MIC_50_) assay

The MIC_50_ assay was conducted according to the method described by Choo et al. [40], with modifications. Briefly, a bacterial inoculum (200 μL of a culture containing 1.5×10^6^ CFU/mL) was dispensed into each well of a 96-well microtiter plate containing serial dilutions of each recombinant protein. Phosphate-buffered saline (PBS) was used as a negative control. The plates were incubated at 28L°C for 24 hours, and bacterial growth was assessed by measuring OD at 600 nm. The MIC_50_ was defined as the lowest protein concentration that resulted in at least 50% inhibition of bacterial growth. Each assay was performed in duplicate for every bacterial strain.

### Bacterial binding assay

The binding of rAtKPI-1T, rTgPI-1, rTgPI-1_NT_, or rTgPI-1_CT_ to microorganisms was performed according to previous methods [41,42]. Briefly, a total of 200 µg of each recombinant inhibitor was incubated with 1 × 10L CFU of bacteria in 1 mL of tris-buffered saline (TBS, 100 mM Tris-HCl, pH 8.0, 10 mM NaCl) under gentle rotation at 28L°C for 1 hour. Following incubation, cells were pelleted by centrifugation and washed three times with TBS to remove unbound protein. Bound proteins were then eluted by resuspending the pellet in 100 µL of TBS containing 12% SDS and agitating for 10–15 minutes. The microorganisms were subsequently washed twice with 1 mL of TBS. Both the SDS-eluted fraction and the final washed pellet were analyzed by 15% SDS-PAGE. Protein detection was performed via Western blot using polyclonal antibodies against either rAtKPI-1T or rTgPI-1. Proteins recovered in the SDS-eluted fraction were classified as weakly bound, while those remaining associated with the pellet were considered tightly bound.

### Indirect Immunofluorescence assay

Overnight bacterial cultures were diluted 1:100 in LB medium and grown until reaching an ODLLL of approximately 0.6. Cells were harvested by centrifugation at 4000 × g for 5 minutes and washed twice with PBS. The resulting pellet was resuspended in 1 mL of PBS and incubated with 20 µg of recombinant protein inhibitors for 1 hour at 28°C with continuous shaking. For fixation, cells were resuspended in 1 mL of 4% formaldehyde prepared in PBS and incubated at room temperature for 30 minutes. After fixation, samples were washed with PBS and blocked with PBS containing 2% bovine serum albumin (BSA) for 20 minutes at room temperature. Primary antibody incubation was carried out overnight using polyclonal antibodies specific to rTgPI-1 or rAtKPI-1T (dilution 1:200). After washing three times with PBS (5 minutes each), cells were incubated for 1 hour with Alexa Fluor 488-conjugated goat anti-mouse IgG secondary antibody (1:4000; Invitrogen), followed by an additional three washes with PBS. Chromosomal DNA was stained with 2.8 mM 4′,6-diamidino-2-phenylindole (DAPI; Molecular Probes) for 10 minutes. Samples were mounted and visualized using a Carl ZEISS Axio Observer 7 inverted microscope, featuring Colibri 7 illumination and both Axiocam 305 color and Axiocam 503 mono cameras. Carl ZEISS ZEN 2.3 (Blue edition) software was used for image acquisition and processing. Fluorescence signals corresponding to DAPI (blue) and Alexa Fluor 488 (green) were captured in separate channels and merged using Image-Pro Plus software (version 4.5, Media Cybernetics).

### Conidial germination assay

The analysis of the antifungal capacity of rTgPI-1_NT_ and rTgPI-1_CT_ on the germination of *B. cinerea* conidia was performed as previously described [30]. Recombinant inhibitors were prepared at final concentrations of 286 nM or 83 nM in 10 mM Tris-HCl and combined with 20 mL of a conidial suspension containing 1.5 × 10^4^ conidia/mL in 25 mM potassium phosphate buffer (pH 5.0) supplemented with 2 M glucose. The mixtures were incubated at 25 °C in individual wells of a 96-well plate (Immuno Plate Maxisorp; Nunc, New York, USA). Conidia germination was assessed microscopically (Nikon Diaphot-TMD, Tokyo, Japan) after 3, 6, and 9 hours. Conidia were considered as germinated if the germ tube exceeded twice the largest diameter of the spore body. For each well, more than 100 conidia were counted, and four technical replicates were analyzed per condition. The entire assay was repeated in three independent experiments. As a negative control, eluates from non-transformed *E. coli* BL21 (DE3) or *E. coli* M15, dissolved in 10 mM Tris-HCl, were used.

## Results

### Expression and purification of the rAtKPI-1T, rTgPI-1, and two truncated versions of rTgPI-1 (rTgPI-1_NT_ and rTgPI-1_CT_)

The recombinant KPIs (rKPIs) were expressed as soluble proteins fused to a His-tag at the N-terminus. The rAtKPI-1T was expressed in *E. coli* BL21 (DE3) as previously described [30], while rTgPI-1, rTgPI-1_NT,_ and rTgPI-1_CT_ were expressed in *E. coli* M15 as previously described [37,38]. The truncated versions of rTgPI-1 have two Kazal domains each, with affinity for trypsin and chymotrypsin for rTgPI-1_NT,_ and with affinity for elastase and chymotrypsin for rTgPI-1_CT_ [37]. After 0.5 mM IPTG induction, the rKPIs were purified with Ni-NTA affinity chromatography from precipitates obtained from *E. coli*. SDS-PAGE analysis showed that rKPIs were conveniently purified using the eluent from the 250 mM imidazole wash (Fig. 1S).

### Bacterial growth inhibition activity of the rAtKPI-1T, rTgPI-1, rTgPI-1_NT_, and rTgPI-1_CT_

Because rAtKPI-1T and rTgPI-1 showed a strong antifungal activity, inhibiting the germination rate of *B. cinerea* conidia *in vitro* [30], we now ask whether these rKPIs could also exhibit a wide antimicrobial activity, including against plant pathogenic bacteria. Therefore, we examined the inhibitory capacity of the rAtKPI-1T and rTgPI-1 against *P. syringae* DC 3000, *P. syringae* (AvRpm1) and *P. viridiflava in vitro*. We also analyzed two truncated versions of rTgPI-1 (rTgPI-1_NT_ and rTgPI-1_CT_), given that these versions were previously shown to have different specificities [37]. Bacterial cultures were grown in the presence of each rKPI and growth was monitored spectrophotometrically (Fig. 1). The four rKPIs inhibited growth of the three bacterial species when were used at a concentration of 7 μM (Fig. 1A). However, for the concentration of 3.5 μM, significant differences in the inhibitory capacity between the rKPIs were observed for the three bacterial species (Fig. 1B). Therefore, taking into account this result, the Minimum Inhibitory Concentration (MIC_50_) was determined using concentrations from 1 μM to 10 μM of each rKPI.

**Fig. 1.**
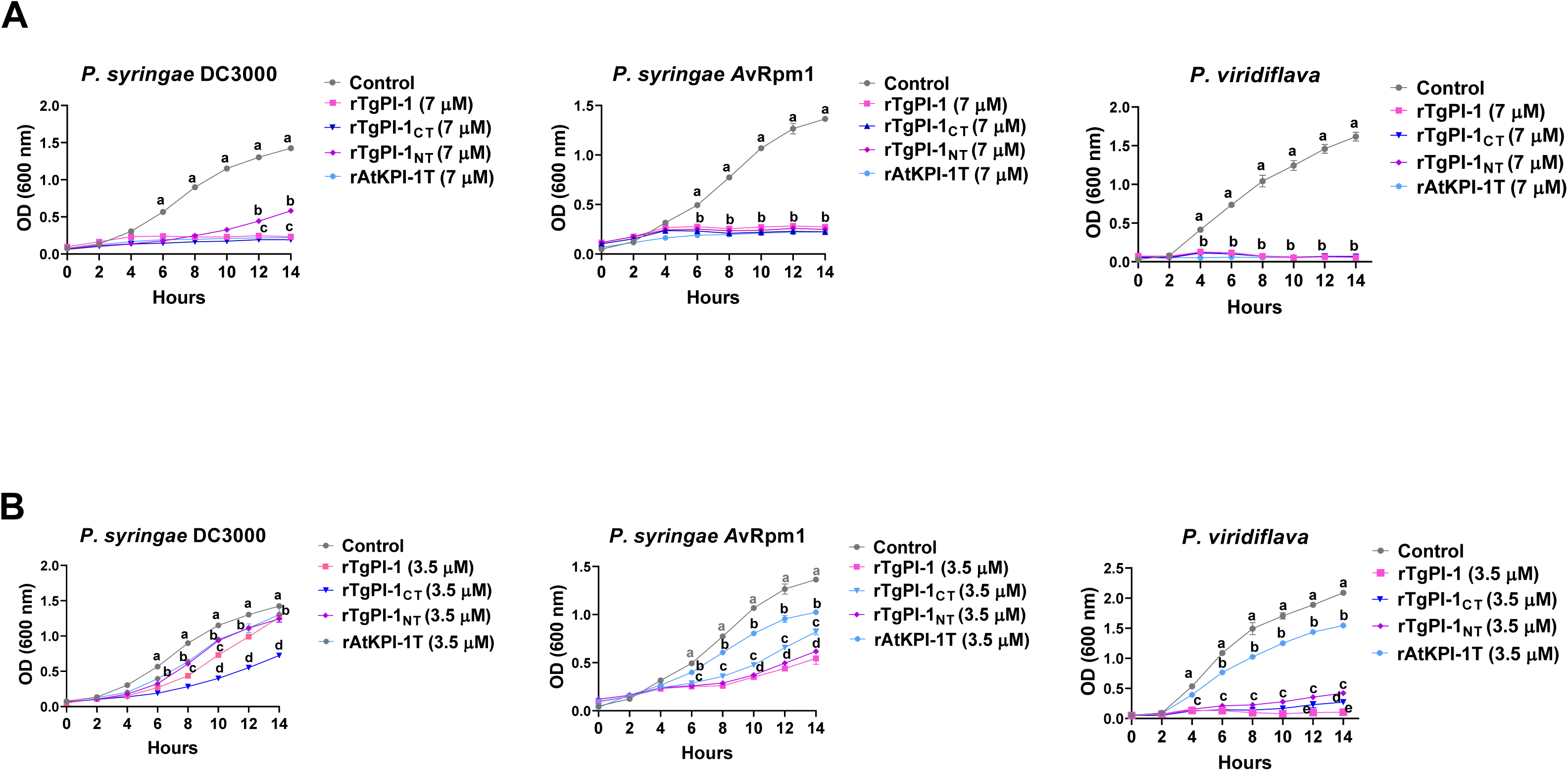
Antibacterial activity of rAtKPI-1T, rTgPI-1, rTgPI-1_NT_, and rTgPI-1_CT_ on *P. syringae* DC3000, *P. syringae* (AvRpm1), and *P. viridiflava*. The cultures were grown with 7 µM (A) or 3.5 µM (B) of rKPIs or PBS as a negative control. Error bars represent the mean ± S.D. (n = 2). Tukey’s multiple comparisons test revealed that there were significant differences between the rKPIs among themselves and with the control group. Different letters represent statistically significant differences (p < 0.05).

Table 1 shows the MIC_50_ values for each rKPI on the different bacterial strains tested. The MIC_50_ values obtained for rTgPI-1, rTgPI-1_NT_, and rTgPI-1_CT_ were lower than those observed for rAtKPI-1T for the three bacterial species. In addition, rTgPI-1, rTgPI-1_NT_, and rTgPI-1_CT_ presented higher MIC_50_ values for *P. syringae* (AvRpm1) than for *P. syringae* DC3000 (Table 1). It is worth mentioning that rTgPI-1 and rTgPI-1_CT_ showed the lowest MIC_50_ values for the three bacterial strains (Table 1), suggesting that the Kazal domains contained in rTgPI-1_CT_ would contribute mainly to the antibacterial activity. However, rTgPI-1_NT_ also presents a strong antibacterial effect. This indicates that both contribute to the recognition of separate targets, and their combination could be more efficient than the recognition of one alone. Therefore, we analyzed whether a synergistic or additive effect between both truncated versions of rTgPI-1 could occur. A bacterial growth inhibition assay was carried out where each bacterial strain was grown in the presence of rTgPI-1, rTgPI-1_NT_, rTgPI-1_CT_, or an equimolar mixture of rTgPI-1_NT_ and rTgPI-1_CT_ (Fig. 2). For both *P. syringae* strains, we did not observe a synergistic or additive effect of the truncated versions of rTgPI-1. However, for *P. viridiflava*, we observed a similar reduction in bacterial growth between bacteria grown in the presence of the mixture of rTgPI-1_NT_ + rTgPI-1_CT_ and those grown in the presence of rTgPI-1, suggesting an additive effect between the four Kazal domains contained in rTgPI-1 (Fig. 2).

**Fig. 2.**
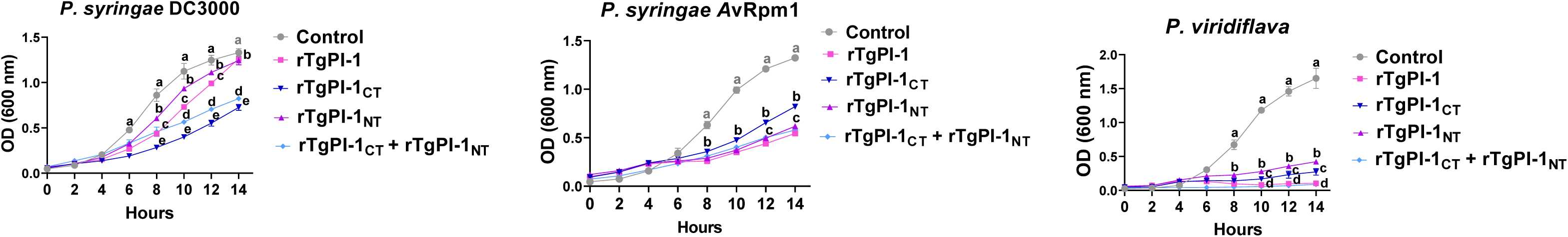
Additive effect on *P. syringae* DC3000, *P. syringae* (AvRpm1), and *P. viridiflava* growth of both truncated versions of rTgPI-1. Each of the bacterial strains was grown in the presence of 3.5 µM of rTgPI-1, rTgPI-1_NT_, rTgPI-1_CT_, or an equimolar mixture of rTgPI-1_NT_ + rTgPI-1_CT_. Error bars represent the mean ± S.D. (n = 2). Tukey’s multiple comparisons test revealed that there were significant differences between the rKPIs among themselves and with the control group. Different letters represent statistically significant differences (p < 0.05).

**Table 1.** Antibacterial activity of recombinant Kazal-type protease inhibitors, expressed as MIC□□ (μM), against *Pseudomonas* strains.

| Bacteria | MIC<sub>50</sub> (μM) |  |  |  |
| --- | --- | --- | --- | --- |
|  | rTgPI-1 | rTgPI-1<sub>CT</sub> | rTgPI-1<sub>NT</sub> | rAtKPI-1T |
| *P. syringae* DC3000 | >2 | >3 | >4 | >10 |
| *P. syringae* (AvRpm1) | >4 | >4 | >6 | >8 |
| *P. viridiflava* | >1 | >1 | >2 | >10 |

### Bacterial binding activity of the rAtKPI-1T, rTgPI-1, rTgPI-1_NT_, and rTgPI-1_CT_

A binding assay was performed to analyze whether rKPIs could bind to microorganisms, and in the case of rTgPI-1, which of the Kazal domains mainly contributed to the binding activity. To answer this question, the truncated versions of rTgPI-1 were evaluated. The result showed that rAtKPI-1, rTgPI-1, and rTgPI-1_NT_ could bind to the three *Pseudomonas* species (Fig. 3A). rAtKPI-1T was the only protein detected in the pellet of the preparations after treatment with SDS, and rTgPI-1 and rTgPI-1_NT_ were detected only in the supernatant (Fig. 3A). rTgPI-1_CT_ was not detected in either the pellet or the supernatant after treatment with SDS (Fig. 3A). Based on this result, the step in the procedure in which rTgPI-1_CT_ was retained was analyzed by Western blot, and we observed that it was detected in the step before treatment with SDS (Fig. 3B).

**Fig. 3.**
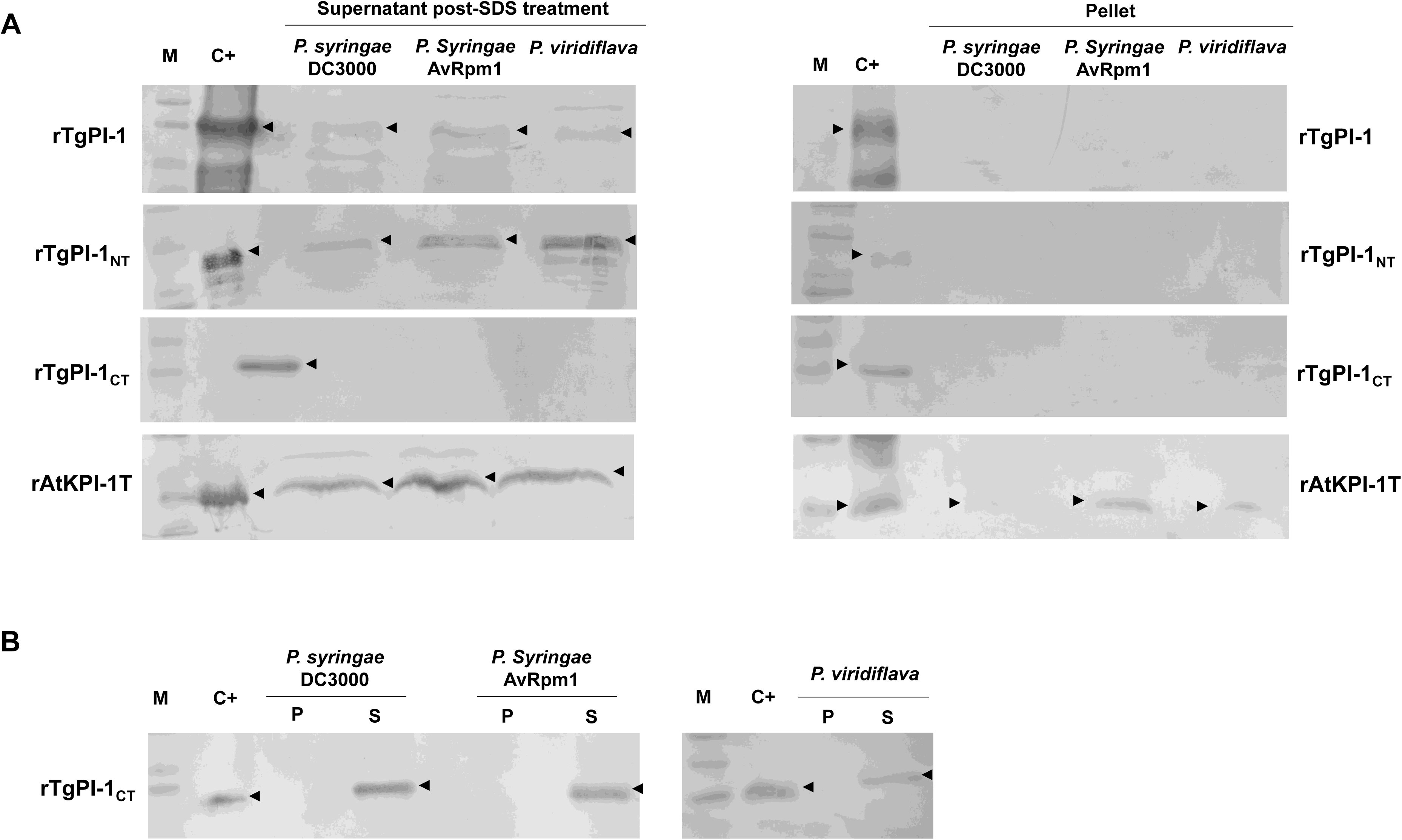
Microbial binding of rAtKPI-1T, rTgPI-1, rTgPI-1_NT_, and rTgPI-1_CT_ to *P. syringae* DC3000, *P. syringae* (AvRpm1), and *P. viridiflava*. rKPIs in the SDS treatment from microorganisms and the pellets after SDS treatment were detected by Western blot with anti-rAtKPI-1 or anti-rTgPI-1 polyclonal antibodies, and the respective rKPIs were used as positive controls. (A) Microbial binding activity of rKPIs in the supernatant post-SDS treatment and in the washed pellet after SDS elution (Pellet). (B) Microbial binding activity of rTgPI-1_CT_ in the supernatant (S) and pellet (P) pre-SDS treatment. M: molecular weight marker for pre-stained proteins. C+: purified rKPIs.

Since the results obtained by Western blot suggest the rKPIs have differences in bacterial binding activity, we performed an indirect immunofluorescence staining (IIF) to confirm the direct binding of the rKPIs to bacteria. The results showed that rTgPI-1 was localized in the cell walls of the three *Pseudomonas* strains (Fig. 4), whereas the truncated versions of rTgPI-1 showed a different pattern of binding. rTgPI-1_NT_ was localized in the cell surface of *P. syringae* DC3000 and AvRpm1, whereas rTgPI-1_CT_ was localized in *P. syringae* AvRpm1 and *P. viridiflava* (Fig. 4). rAtKPI-1T showed a similar affinity to rTgPI-1_NT_, localizing in *P. syringae* DC3000 and AvRpm1 (Fig. 4).

**Fig. 4.**
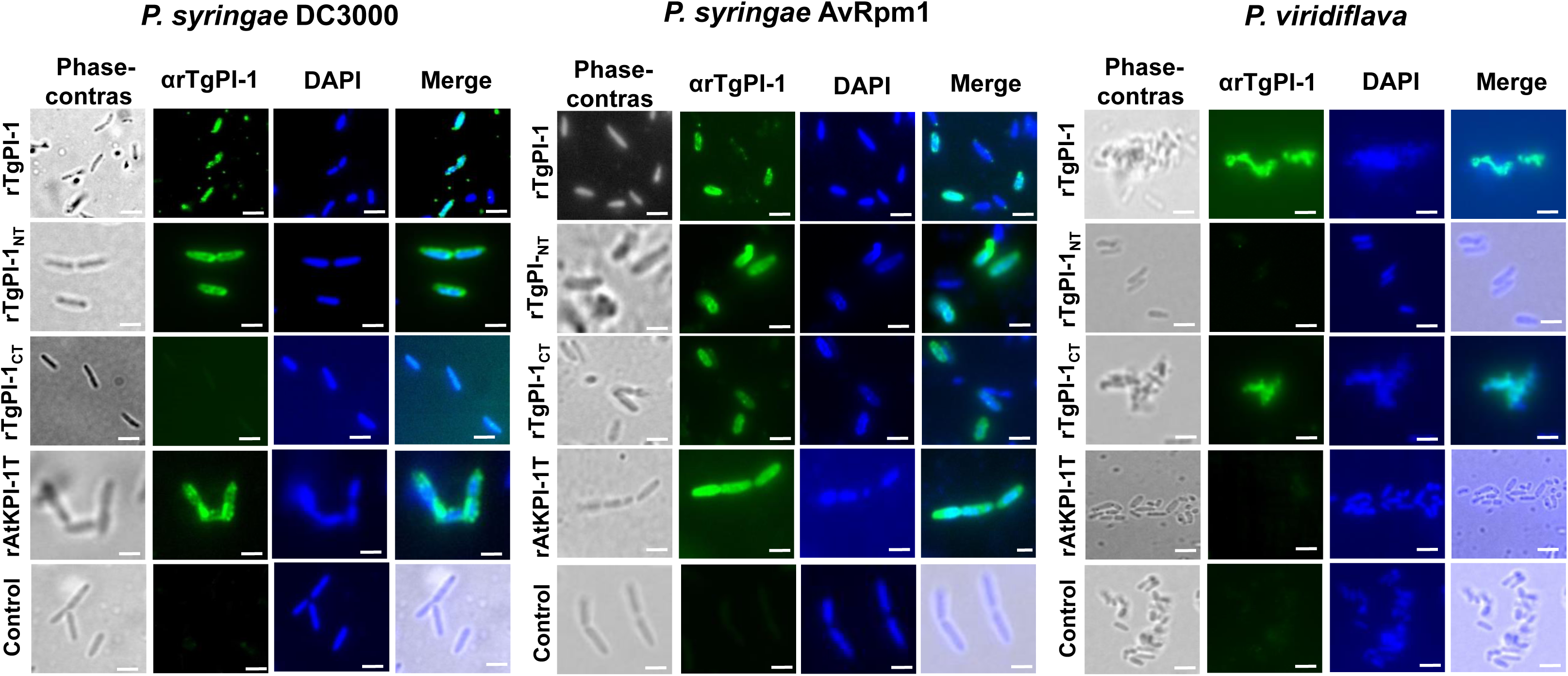
Direct binding activity of rAtKPI-1T, rTgPI-1, rTgPI-1_NT_, and rTgPI-1_CT_ to bacteria. Immunofluorescence staining was performed to visualize the binding of rKPIs to living *P. syringae* DC3000, *P. syringae* (AvRpm1), and *P. viridiflava*. Bacteria were incubated with mouse anti-rTgPI-1 or anti-rAtKPI-1T polyclonal antibody as primary antibody and Alexa Fluor 488 goat anti-mouse IgG (green color) as secondary antibody. Chromosomic DNA were stained with DAPI. The three images were merged (phase contrast, rKPIs+, chDNA). Scale bar represents 1 µm. This image is representative of a larger field of view, and data are from a representative experiment performed three times. Phase contrast, green and blue fluorescence were recorded separately, and the images were merged using image-pro plus 4.5.

### *In vitro* inhibition of *B. cinerea* conidia germination by rTgPI-1_NT_ and rTgPI-1_CT_

In a previous study, we evaluated the inhibition capacity of rAtKPI-1T and rTgPI-1 on the germination of *B. cinerea* conidia [30]. Here, we analyzed the germination of *B. cinerea* conidia in culture medium supplemented with rTgPI-1_NT_ and rTgPI-1_CT_ to determine whether the truncated versions of rTgPI-1 can exert antifungal effects.

Conidial germination of *B. cinerea* was markedly reduced after 6 and 9 hours of incubation in media supplemented with rTgPI-1 truncated versions, compared to the control lacking such inhibitors (Fig. 5A). Notably, rTgPI-1_CT_ exhibited a significantly stronger inhibitory effect than rTgPI-1_NT_ at a concentration of 286 nM (Fig. 5A). In addition, microscopic examination revealed that rTgPI-1_CT_ and rTgPI-1_NT_ treatments impaired germ-tube elongation, in contrast to the untreated control, which displayed typical germ-tube development (Fig. 5B).

**Fig. 5.**
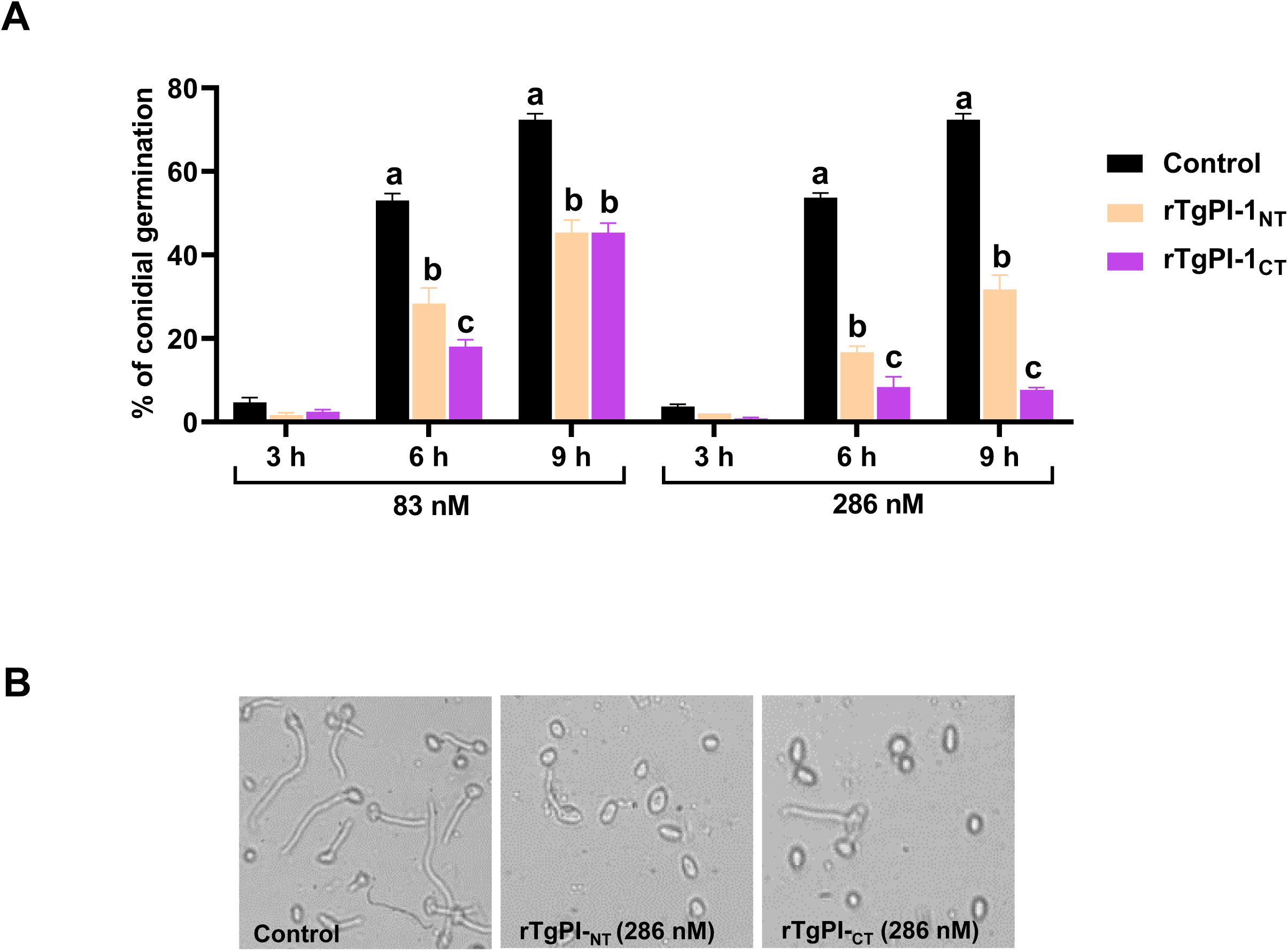
Germination of *Botrytis cinerea* in medium supplemented with rTgPI-1_NT_ and rTgPI-1_CT_. (A) Inhibition of conidial germination. Results are means of four replicates ± S.D. and are representative of three experiments. Statistical analysis was performed by one-way ANOVA using Bonferroni’s Multiple Comparison Test. Comparisons are valid only within each time period. Different letters represent statistically significant differences (p < 0.05). (B) Germ-tube growth of *B. cinerea* conidia after incubation for 9 h *in vitro* with 286 nM of rKPIs. Control: addition of an eluate from *E. coli* M15; rTgPI-1_NT_: addition of 83 or 286 nM of rTgPI-1_NT_; rTgPI-1_CT_: addition of 83 or 286 nM of rTgPI-1_CT_.

## Discussion

Plant pathogens have evolved mechanisms to evade host detection and modulate host signaling pathways to their advantage. Proteases significantly contribute to these infection-associated processes [43]. Thus, several protease inhibitors are likely involved in protecting the host from microbial proteases and may participate in the immune defense response. Generally, the protease inhibitors are one of the first proteins to be activated in plants as a defense mechanism against pathogens, promoting resistance [29,44,45]. In this sense, it is relevant to study the mechanisms of action of these inhibitors to advance in the design of biotechnological tools for the control plant diseases. Although KPIs have been isolated from a wide variety of organisms and their role in the modulation of the innate immune response and their antimicrobial activity have been reported in various studies, few KPIs from plants have been functionally characterized [29,30,45]. In this study, our results demonstrate that rKPIs exhibit antimicrobial activity against phytopathogenic bacteria and fungi. Specifically, rAtKPI-1T and rTgPI-1, previously reported as fungal growth inhibitors in *B. cinerea* [30], also exhibited antibacterial effects against *P. syringae* DC3000, *P. syringae* (AvRpm1), and *P. viridiflava*. The inhibitory effects were concentration-dependent, with significant differences among KPIs observed at 3.5 μM, allowing the determination of MIC_50_ values. The lower MIC_50_ values for rTgPI-1 and its truncated versions (rTgPI-1_NT_ and rTgPI-1_CT_) compared to rAtKPI-1T demonstrate a higher antibacterial activity. In this regard, some reports show that KPIs can interfere with bacterial virulence by specifically targeting essential serine proteases involved in pathogenesis, suggesting that the observed antimicrobial effect could be related to the type of serine proteases inhibited by each rKPI [24,25]. Since rAtKPI-1T has been found to specifically inhibit elastase and subtilisin [30], whereas rTgPI-1 has a broader inhibition spectrum, targeting trypsin, chymotrypsin, and elastase [20], it is plausible that the inhibitory activity of these rKPIs affects bacterial fitness and pathogenicity differently depending on their protease targets. In particular, the lower MIC_50_ observed for rTgPI-1_CT_ suggests that the Kazal-type domains in this variant are particularly effective in inhibiting key proteases essential for bacterial growth. These results are consistent with previous studies that described how serine protease inhibition can reduce bacterial virulence [29].

Several studies have shown that other families of plant SPIs exhibit microorganism-binding properties [46]. Certain Kunitz-type inhibitors have demonstrated a strong affinity for the surface of Gram-negative and Gram-positive bacteria, modulating their growth and virulence [47–49]. Additionally, some SPIs have been reported to act by binding to bacterial lipopolysaccharides, altering cell membrane stability, and affecting bacterial structural integrity [50,51]. These findings reinforce the idea that bacterial surface binding may be a key mechanism in the antibacterial activity of the rKPIs studied here. In fact, the bacterial binding assays provided further insights into the mode of action of these inhibitors. The ability of rAtKPI-1T, rTgPI-1, and rTgPI-1_NT_ to bind to the three *Pseudomonas* species suggests a role in direct bacterial interaction. In addition, the detection of rAtKPI-1T in the bacterial pellet after SDS treatment indicates strong and possibly irreversible binding to bacterial components [52]. In contrast, the absence of rTgPI-1_CT_ in either the pellet or supernatant post-SDS treatment, but its detection before treatment, suggests a weaker or transient interaction with the bacterial surface [35,52]. The indirect immunofluorescence staining (IIF) also showed differences in bacterial binding affinities among the rKPIs. rTgPI-1 localized in the cell surface of all three *Pseudomonas* species, while the truncated versions exhibited strain and species-specific affinities. rTgPI-1_NT_ showed affinity for *P. syringae* DC3000 and *P. syringae* (AvRpm1), whereas rTgPI-1_CT_ was associated with *P. syringae* (AvRpm1) and *P. viridiflava*. This differential binding pattern is consistent with other works on plant SPIs [34,53]. A study on a Kunitz-type inhibitor from *Inga laurina* showed different interactions with bacterial pathogens depending on the composition of the bacterial cell membrane [34]. Similarly, certain KPIs derived from organisms other than plants, have been reported to recognize specific bacterial structures through interactions with outer membrane components [42,52]. The binding pattern of rAtKPI-1T, similar to that of rTgPI-1_NT_, suggests functional similarities between these proteins, possibly reflecting conserved mechanisms of bacterial recognition and interaction [30,37]. Given that Kazal-type domains can determine pathogen-binding specificity, the results obtained here reinforce the notion that the molecular architecture of these inhibitors influences their antimicrobial activity.

Interestingly, some KPIs have also been shown to display complex and sometimes inconsistent relationships between binding, protease inhibition, and microbial growth suppression. rCqKPI from *Cherax quadricarinatus* was found to selectively bind a wide range of microorganisms, including Gram-positive and Gram-negative bacteria as well as fungi, and partially inactivate bacterial secretory proteases [35]. Notably, this KPI exhibited a strong inhibitory activity against *Candida albicans* proteases and growth, despite weak binding, and an opposite profile against *Saccharomyces aureus* (strong binding but no growth inhibition). Similar discrepancies have been described for other crustacean KPIs such as hcPcSPI2 from *Procambarus clarkii*, which strongly bound to *E. coli* and *S. aureus* without affecting their growth [54], and SPIPm4/SPIPm5 from *Penaeus monodon*, which inhibited subtilisin activity but lacked bacteriostatic effects [55]. The distinct binding and antimicrobial profiles observed for rTgPI-1_NT_ and rTgPI-1_CT_ support this broader concept that KPIs can differentially affect microbial physiology depending on their target proteases and modes of interaction. These findings highlight that the bacteriostatic action of KPIs not only depends on the protease inhibition potency, but also on the specific role of the target serine protease in microbial physiology. These findings suggest that the influence of KPIs on bacteria is multifaceted, and their growth inhibition capacity reflects a cumulative effect of multiple interactions [35,54,55].

In addition to their antibacterial effects, rTgPI-1_NT_ and rTgPI-1_CT_ also exhibited antifungal activity against *B. cinerea* conidia. Both truncated proteins significantly reduced conidial germination after 6 and 9 hours of incubation, with rTgPI-1_CT_ showing a markedly stronger inhibitory effect than rTgPI-1_NT_. Microscopic observations confirmed that these treatments impaired germ-tube elongation compared to untreated controls. These findings indicate that truncated rKPIs retain antifungal capacity observed for rTgPI-1 [30], suggesting that specific Kazal-type domains, particularly those in rTgPI-1_CT_, play a crucial role in inhibiting early infection processes of necrotrophic fungi. This is in agreement with previous reports highlighting the involvement of SPIs in plant defense against fungal pathogens [29], and extends the understanding of how individual Kazal-type domains contribute to antifungal activity. Importantly, our results demonstrate that rTgPI-1_CT_ exerts a much stronger suppression of *B. cinerea* germination than rTgPI-1_NT_, suggesting that the C-terminal domains of TgPI-1, which contain Leu at the P1 site, may preferentially inhibit elastase-like fungal proteases essential for early infection. Conversely, the weaker inhibition by TgPI-1_NT_ could be related to Arg at the P1, which is more typical of trypsin inhibition, indicating that different domains of the same inhibitor can contribute unequally to antifungal defense [37]. This functional divergence between the N-terminal and C-terminal domains mirrors the domain-specific specificity described in other KPIs, providing a possible mechanistic explanation for the differential inhibition observed. Therefore, the antifungal activity of the truncated versions of TgPI-1 not only validates the role of serine protease inhibition in limiting *B. cinerea* infection but also underscores the relevance of domain composition in shaping the spectrum and intensity of the antifungal response. In addition, previous studies demonstrated that rAtKPI-1T strongly inhibited elastase and subtilisin, while showing only weak activity against trypsin. It was associated with significant suppression of *B. cinerea* conidial germination and germ-tube elongation [30]. Therefore, the biochemical specificity of rAtKPI-1T provides mechanistic support for the antifungal effects observed in our assays, suggesting that the inhibitory activity against subtilisin- and elastase-like proteases may directly impair fungal infection structures. Moreover, the structural determinants of this specificity, such as the composition of the reactive loop and the identity of the P1 residue, may play a critical role in defining the inhibitory spectrum of Kazal-type domains. This highlights the importance of combining structural analyses with functional assays to understand how individual domains contribute to antifungal defense.

## Conclusion

Overall, our findings provide evidence of the antimicrobial potential of rKPIs mediated by their ability to bind microbial surfaces and inhibit essential proteases. The observed differences in MIC_50_ values, binding affinities, and localization patterns suggest functional diversity among the Kazal domains of these SPIs. In particular, the contrasting behaviors of rTgPI-1_NT_ and rTgPI-1_CT_ highlight the value of domain truncations as experimental tools to dissect the contribution of individual Kazal modules to antimicrobial activity. This domain-level analysis provides novel insights into how specific structural elements shape both antibacterial and antifungal responses. These results expand on previous studies [30] and highlight the need for further structural and functional analyses to elucidate their mechanism of action. Beyond their relevance for basic research, the demonstration that plant KPI, and particularly the truncated variants of TgPI-1, retain strong and selective antimicrobial activities suggests that these proteins could be rationally engineered as novel biocontrol agents. Such strategies may help develop sustainable approaches for crop protection, reducing reliance on chemical pesticides and enhancing plant resistance against phytopathogens. Future research should focus on characterizing the molecular interactions between KPIs and their microbial targets to optimize their potential application in crop protection strategies.

## Acknowledgements

M. Laguía-Becher, V. A. Sander, F. L. Pieckenstain, and M. Clemente are members of Consejo Nacional de Investigaciones Científicas y Técnicas (CONICET). V. A. Sander, F. L. Piechenstain, and M. Clemente are professors at Universidad Nacional General San Martín (UNSAM). M. Laguía-Becher is head of practical work at UNSAM. M. A. Sánchez is a doctoral fellow of Fondo para la Investigación Científica y Tecnológica (FONCyT) and a practical work assistant at UNSAM. K. N. Strack is a post-doctoral fellow of CONICET and a practical work assistant at UNSAM. L. F. Mendoza-Morales was a doctoral fellow of CONICET. Currently is a professor at Universidad Industrial de Santander, Málaga, Colombia. V. A. Ramos-Duarte was a doctoral fellow of CONICET. Currently is a technical manager at the Technical University of Munich, Germany. The authors thank Patricia Uchiya (CIC-pba), José Luis Burgos (CONICET) for their technical assistance, and Carlos Alberici (CONICET), Gustavo Muñoz (CONICET), and Claudio Guida (CONICET) for their maintenance of research instruments.

## Author contributions

M.A.S.: Formal analysis, Investigation, Methodology, Visualization, Writing – review & editing. K.N.S., L.F.M.M., V.A.R.D., and E.P.: Methodology, Visualization, Writing – review & editing. V.A.S., M.L.B., and F.L.P.: Conceptualization, Writing – review & editing. M.C.: Conceptualization, Funding acquisition, Project administration, Supervision, Writing – original draft.

## Data availability

The datasets used and/or analyzed during the current study are available from the corresponding author on reasonable request.

## Supplementary Figures

**Fig. S1.**
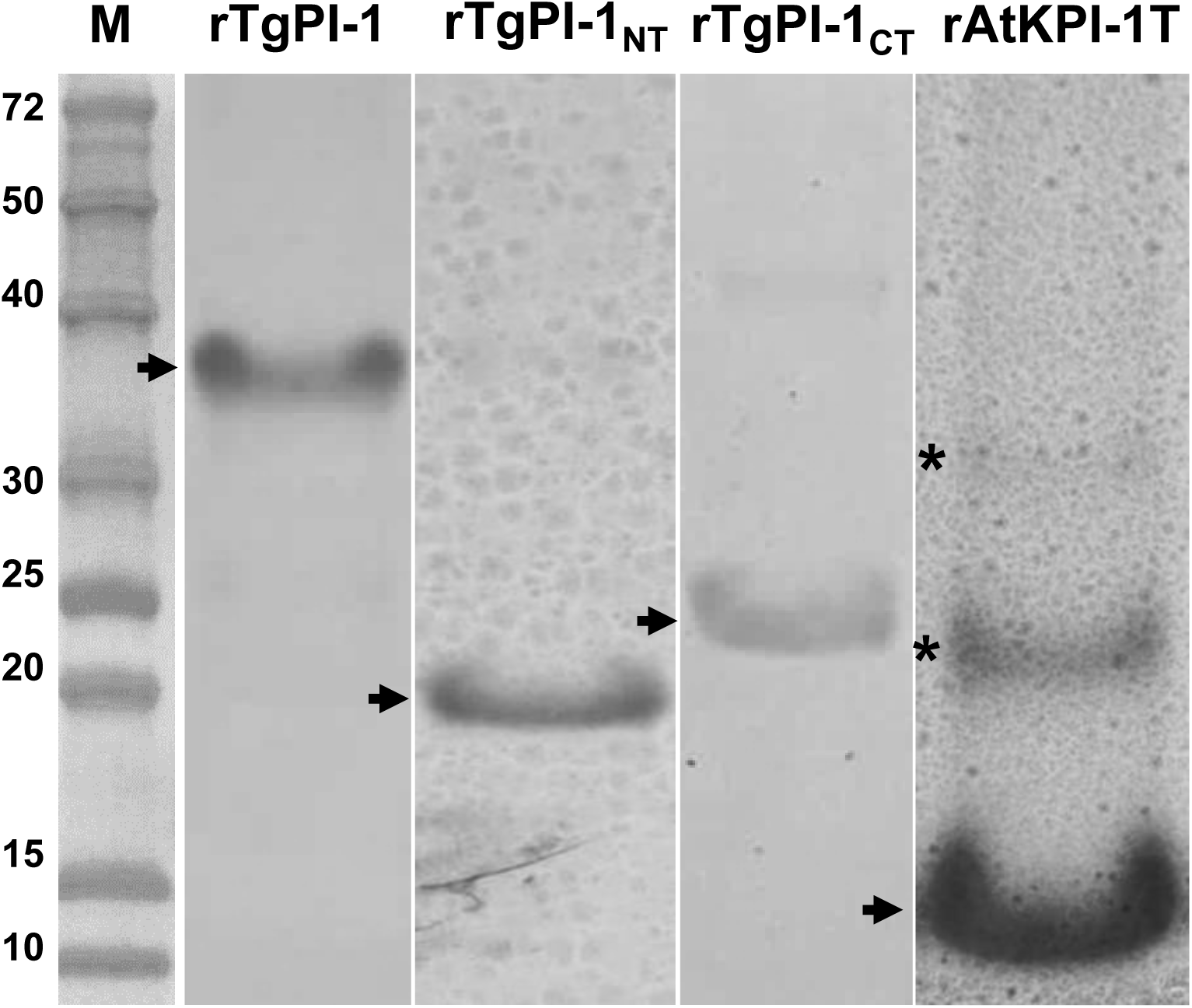
Expression and purification of the recombinant Kazal-type protease inhibitors (rKPIs). Purification of rAtKPI-1T, rTgPI-1, rTgPI-1_NT_, and rTgPI-1_CT_ was checked in a Coomassie blue-stained polyacrylamide gel. rKPIs were eluted using a Ni+ affinity chromatography column.

